# Local selection shaped the diversity of European maize landraces

**DOI:** 10.1101/2024.06.07.597898

**Authors:** Margarita Takou, Kerstin Schulz, Markus G Stetter

## Abstract

The introduction of populations to novel environments can lead to a loss of genetic diversity and the accumulation of deleterious mutations due to selection and demographic changes. We investigate how the recent introduction of maize to Europe shaped the genetic diversity and differentiation of European traditional maize populations and quantify the impact of its recent range expansion and consecutive breeding on the accumulation of genetic load. We use genome-wide genetic markers of almost 2,000 individuals from 38 landraces, 155 elite breeding lines and a large set of doubled haploid lines derived from two landraces to find extensive population structure within European maize, with landraces being highly differentiated even over short geographic distances. Yet, diversity change does not follow the continuous pattern of range expansions. Landraces maintain high genetic diversity that is distinct between populations and does not decrease along the possible expansion routes. Signals of positive selection in European landraces that overlap with selection in Asian maize suggest convergent selection during maize introductions. At the same time, environmental factors partially explain genetic differences across Europe. Consistent with the maintenance of high diversity, we find no evidence of genetic load accumulating along the maize introduction route in European maize. However, modern breeding likely purged highly deleterious alleles but accumulated genetic load in elite germplasm. Our results reconstruct the history of maize in Europe and show that landraces have maintained high genetic diversity that could reduce genetic load in the European maize breeding pools.

## Introduction

Species distributions are the result of range expansion to new environments, such as the post glacial colonization of Northern Europe (Hewitt 2000). These dynamic population genetic processes have a strong influence on the genetic diversity of the expanding species (Excoffier et al. 2009), including a decrease in the genetic diversity due to increased drift and the subsequent genetic differentiation among the newly established and core populations (Austerlitz et al. 1997; Excoffier et al. 2009; Slatkin and Excoffier 2012; Wang et al. 2017; Takou et al. 2021). At the same time, deleterious mutations accumulate at the front of the expansion range creating the genetic load (de Pedro et al. 2021; González-Martínez et al. 2017; Peischl et al. 2013). As locally adapted populations move to novel environments (Colautti and Barrett 2013; Savolainen et al. 2013), the adaptive potential of populations at the range edges can be compromised (Excoffier et al. 2009). Rapid range expansions are expected to become more frequent as climate change alters the natural environments, which will push species to new suitable conditions outside their original range (Waldvogel et al. 2020). Crops have spread rapidly around the globe and likely experienced the effects of range expansion (Huang et al. 2022). They could have encountered novel selective pressures caused by the different climate, soil compositions, and ecological dynamics of the new regions, as well as the human-driven selection for agricultural traits (Purugganan and Fuller 2009; Purugganan 2019). This interplay between natural selection and human-driven intervention ultimately resulted in the appearance of local traditional crop varieties (Huang *et al*. 2022).

One of the most important crops worldwide is maize, which has shown great adaptability to locations outside its initial range with one of the broadest cultivated ranges of all crops today (Tenaillon and Charcosset 2011). Maize domestication started approximately 9,000 years ago in a small region in Mexico, where the wild grass species teosinte, (*Zea mays ssp parviglumis* and *Zea mays ssp mexicana*), have given rise to the crop we know today (Tenaillon and Charcosset 2011; Yang *et al*. 2017b, 2023). Over time, maize spread throughout the Americas, facilitated by human movement (Kistler *et al*. 2018). The early dispersion process was characterized by two routes: one northward to the USA and Canada, and another extending southward to South America and the Caribbean coast (Tenaillon and Charcosset 2011; Kistler *et al*. 2018). This human mediated range expansion has led to decreased genetic diversity, introgression events and local adaptation across the colonization route (Wang *et al*. 2017; Hufford *et al*. 2012; Arca *et al*. 2023), which creates the opportunity to study range expansion and the dynamics of selection and decreasing genetic diversity. Hence, crops such as maize might have experienced similar range expansion forces that are predicted for natural populations.

European maize landraces offer a compelling system to explore the recent introduction and expansion at potential range edges. While extensive research has been done on the evolution, domestication, and traditional use of primary American maize landraces (Hufford et al. 2012; Yang et al. 2023; Wang et al. 2017), the secondary range expansion and development of their European counterparts have been little investigated (Revilla Temiño et al. 2003; Tenaillon and Charcosset 2011). Unlike the gradual expansion outside of the domestication area, maize’s arrival in Europe is characterized by an abrupt introduction to a novel environment around 500 years ago (Janick and Caneva 2005; Brandenburg et al. 2017). Moreover, only a relatively small number of plants were initially brought from the Americas, likely resulting in a strong bottleneck (Tenaillon and Charcosset 2011). After its first introduction into Europe from the Caribbeans through Spain, there is evidence for further diffusion routes via France and Italy to the rest of Europe, as well as a secondary introduction from North America to Central Europe (Brandenburg et al. 2017; Leng et al. 1962; Rebourg et al. 2003). Additionally, the varying local European environments likely required local adaptation, as has been observed in Mexican maize populations (Tittes et al. 2023). Historical accounts state that by the middle of the 16th century maize fields were present across all Europe (Finan 1948; Janick and Caneva 2005; Brandolini and Brandolini 2009), making European maize landraces an invaluable resource for understanding the rapid introduction to new environments.

Here, we study the genetic diversity across Europe and how it was potentially shaped by the multiple introductions of maize to the continent. Using a comprehensive dataset of traditional varieties spanning the range of European maize, we were able to show that European landraces are highly diverse but genetically differentiated. We detect two major clusters of landraces within Europe that might depict their introduction history. In addition, we identify local clusters of similarity between traditional varieties, likely reconstructing historical trade routes. Selection scans in two landrace double haploid (DH) line libraries with almost 1,000 individuals reveal genomic regions that were likely under selection in individual landraces and suggest potential adaptation signatures within Europe. However, European maize does not show signatures of range edge decreased diversity and accumulated genetic load. Together, our results give insights into the history of the rise of European maize in the last centuries.

## Materials and Methods

### Dataset composition

We selected a sample set to represent the majority of maize’s history in Europe. Two primary categories of maize samples were used to form the panel: European landraces and elite lines. The European landraces, which represent authentic European populations (Villa et al. 2005; Casañas et al. 2017), were selected with the aim of capturing a high geographical range. The second category, elite lines, are genetically stable lines that are derived from landrace material (Reif et al. 2005) and were included as points of comparison with European landraces. Furthermore, double haploid lines (DH lines), which are completely homozygous inbred lines of three European landraces (Maqbool *et al*. 2020), were also included.

All data used in the present study are publicly available (Unterseer *et al*. 2016; Mayer *et al*. 2017, 2020). In brief, all individuals used were genotyped at high density with the 600k Affymetrix® Axiom® Maize Array, which is comprised of 616,201 variants, of which 6,759 represent insertions/deletions (Unterseer et al. 2014, 2016). Specifically, 941 DH lines derived from 2 European landraces were taken from Mayer *et al*. (2020), which were divided as follows: 501 DH lines of Kemater Landmais Gelb (KL; Austria) and 409 of Petkuser Ferdinand Rot (PE; Germany). 952 individuals from 35 European maize landraces from Mayer *et al*. (2017). Lastly, 3 landraces from Unterseer *et al*. (2016) and 155 elite lines were added from Unterseer *et al*. (2016). For the following analysis, we clustered the populations in two groups, based on population clustering (Table S1). The final panel had in total 2,954 individuals (Table S1).

### Data Preparation

The merging of the three data sets (Unterseer *et al*. 2016; Mayer *et al*. 2017, 2020) was performed using custom python scripts and all data sets were formatted in the HapMap format using Tassel 5.0 GUI (command: .*/start_tassel*.*pl -Xmx4g*) (Bradbury *et al*. 2007). Post conversion the IUPAC nucleotide codes in the converted files were replaced with the corresponding nucleotide bases as stated in the user guide. For merging, the Unterseer *et al*. (2016) data sets were updated to AGPv4 maize reference genome. All insertions/deletions in the datasets were removed, which resulted in 6,759 and 6,752 markers removed from the Mayer and Unterseer data sets, respectively. For quality control, we checked if the alleles were correctly merged, based on the individual marker information provided by the manufacturer. Additionally, in order to filter out possible sequencing errors and sites with low genotyping quality, we used the Affymetrix Quality classification, and only markers that had quality of “PolyHigh-Resolution”, “MonoHighResolution” or “NoMinorHom” across all datasets were kept, resulting in 419,477 SNPs. As none of the individuals had missing information in more than 0.8% of their sites and the majority of the sites had less than 1% missing information after the above filtering steps, we did not apply any other filtering criteria based on missingness. Subsequently, the dataset tables were merged and adjusted to fit the HapMap format, enabling conversion via TASSEL. Lastly, the Hapmap file was converted back into a common VCF file using TASSEL (Bradbury *et al*. (2007)) The resulting VCF file was then used for the subsequent analysis, each time filtered as indicated within the respective sections.

### Population Structure and Genetic Diversity analysis

To investigate the underlying patterns of variation in this maize dataset a Principal Component Analysis (PCA) was conducted using the scikit allel package (Miles *et al*. 2023) for python v3.7. All individuals of the 35 landraces from Mayer *et al*. (2017), the 3 unique landrace lines from Unterseer *et al*. (2016) AL, FL, and SO, as well as the elite maize lines grouped into dent and flint populations were used for this analysis. We sub-sampled the dataset to exclude the DH lines and reduce the individuals to 1,160 individuals divided into 40 populations, including the dent and flint elite lines, for the PCA. In order to do the PCA, using the scikit allel package, singletons were removed from the sub-sampled dataset and linkage disequilibrium (LD) filtering/pruning was done using 5 iterations, a window size of 500bp, and a step size of 200bp. The PCA was then performed using the function *allel*.*pca()* (Miles *et al*. 2023) and plotted using the matplotlib package (Hunter 2007). We further investigated the clustering of the data by using the software STRUCTURE (Pritchard *et al*. 2000). We run the analysis for up to K=40 and used the cross validation error (cv) to decide the optimal number of clusters.

Using scikit allele package (Miles *et al*. 2023), all pairwise *F*_*ST*_ values between all the 35 European landrace population pairs from Mayer *et al*. (2017) were estimated based on segregating SNPs. We calculated Hudson’s *F*_*ST*_ using the *allel*.*blockwise_hudson_fst()* function with a block size of 100,000 variants (Smaragdov and Kudinov 2020; Bhatia et al. 2013; Hudson et al. 1992). Additionally, we estimated Slatkin’s linearized *F*_*ST*_, as shown in (Bay et al. 2018). To visualize the population differentiation patterns, the calculated *F*_*ST*_ values were plotted with a hierarchical clustered heatmap, using the package ComplexHeatmap in R v4.1.2 (R Core Team 2021; Gu et al. 2016).

We estimated mean expected heterozygosity (*H*_*EXP*_) per population for all landraces to investigate the distribution of genetic diversity and potential patterns of diversity among European landraces across the diverse European landscape. We calculated the expected heterozygosity using the *allel*.*heterozygosity_expected()* function and estimated the mean using the *numpy*.*nanmean()* function (Harris et al. 2020). The geographical map including the *H*_*EXP*_ values was created using the plotly package (Inc. 2015). The sample locations (longitude and latitude values from Mayer *et al*. (2017) were used as proxies for each population’s distribution. We estimated the haversine geographic distance in kilometers of each landrace to the two populations that are approximately the closest to the expected entry points in Europe; the population Tuy (TU; 42.04N, -8.64W) marks the entry point in Spain and the population Barisis (BA; 49.57N, 3.328E) in Central Europe. We estimated all the distances with the function distm of the R package geosphere (Hijmans 2022). For the populations Altreier (AL; Italy), Fleimstal (FL; Italy) and Sornay (SO; France), whose coordinate information was missing, we used the best approximation of their locations. The inbreeding coefficient of each individual was estimated using vcftools (Danecek et al. 2011).

### Spatial gene-flow analysis of population genetic data and selection scans

The Fast Estimation of Effective Migration Surfaces (FEEMS, version 1.0.0) analysis was conducted to investigate non-homogeneous gene flow between the European landraces (Marcus et al. 2021). For this analysis, we used the 35 populations published in Mayer *et al*. (2017), as for the 3 additional landraces in Unterseer *et al*. (2016) we did not have accurate geographic information for them, leaving 834 individuals for 35 landrace populations. Filtering was performed on the final dataset and included the following steps: Non-SNP type SNPs were removed using the command *bcftools view -v snps*, and only biallelic SNPs were selected with *bcftools view -m2 -M2*. As no SNPs with more than 20% missing data were present, we did not remove any sites based on this filter. Monomorphic sites were removed with bcftools (Li 2011) and the input files were prepared with PLINK2 (Chang et al. 2015). We used the FEEMS imputation feature for any present missing data. Specifically, we used scikit’s function SimpleImputer and the imputation strategy of ‘mean’. Additionally, the script was run with the options `scale_snps=True` and `translated=False` and the provided smaller grid size file `grid_100.shp`. A custom smoothing regularization parameter (*λ* = 100) was estimated from a cross-validation procedure using their provided `cross-validation.ipynb`.

### Selection Scans

We searched for signals of selection across the genomes of the DH lines generated for Petkuser Ferdinard Rot (PE DH; Germany) and Kemater Landmais Gelb (KL DH; Austria) as those are naturally phased datasets with approximately 500 individuals each. We used selscan (Szpiech and Hernandez 2014; Szpiech 2024) to estimate the intergrated haplotype score (iHS). For each SNP, we estimated an empirical p-value by dividing its rank of the absolute iHS value, by the number of SNPs. We classified SNPs as being under selection when their p-value was below 0.05 and their absolute iHS value was greater than 2. Then, we extracted either the upstream or downstream genes that were the closest to the selected SNPs. We tested for Gene Ontology enrichment using the online database agriGO v2.2 (Tian *et al*. 2017).

### Climate characterization and climate redundancy analysis

We used historical bioclimatic variables bio1 through bio19 to characterize the population’s environment (Fick and Hijmans 2017). For this analysis, we grouped the populations based on east and west separation into clusters, and we compared the distributions of the environments with a Wilcoxon’s rank test. Moreover, we correlated the bioclimatic variables with the estimated population genetic parameters using Spearman’s *rho* correlation coefficient.

We determined how genetic and environmental factors interact, by performing the multiviarate ordination technique, Redundancy Analysis (RDA) (Capblancq and Forester 2021). We inferred the population structure of the landraces by extracting the first two PCs of a PCA on nearly neutral sites. Near neutral sites were identified as all the 4-fold sites using degenotate from https://github.com/harvardinformatics/degenotate/tree/main. We selected for the climatic variables that fully explain the genetic structure by a forward model building procedure, by using the function rda() from the R package vegan (Oksanen *et al*. 2022). After we identified which bioclimatic variables are part of the model, we tested for the drivers of genetic variation by comparing four different models. A full model that included the selected bioclimatic variables, the first two PCs of neutral sites, latitudinal and longitudinal information as fixed effects; a purely climatic model that incorporated the bioclimatic variables as fixed effects and the geographical and population structure information as ‘condition’; a pure neutral population structure model, which incorporated the PCs as fixed and the rest of the parameters as conditional and finally, a geographic model, which had the longitudinal and latitudinal data as fixed effects and the rest as conditional parameters. We selected for the model that best described the dataset based on their *R*^2^ values via the function RsquareAdj() from the vegan package. All results were plotted with ggplot2 (Wickham 2016).

For each pair of populations we calculated their environmental distance, as described in (Bay *et al*. 2018). We first scaled and centered each bioclimatic variable and subsequently, we calculate the pairwise Euclidean differences between each landrace.

### Genomic load estimations

We estimated genetic load per individual for the landrace populations and elite pools to estimate the accumulation of deleterious mutations. We used the Genomic Evolutionary Rate Profiling (GERP) scores from Kistler *et al*. (2018). To estimate genetic load, for each individual belonging to one of the landraces or elite pools, we scored each site based on the number of derived alleles they have and then summed up the scores across all sites. We excluded 100 individuals that had more than 1.69% of missing sites (95th quantile of the distribution), in order to exclude potential differences in load due to missing information. Furthermore, we estimated the fixed and segregating load within the dent elite and flint breeding pool, as well as for the dent and flint landraces. Finally, we counted the number of highly deleterious alleles, which we defined as sites with GERP score larger than 5, within each sample. We compared the genetic load distributions between the different groups with a Kolmogorov-Smirnov test.

## Results

### Genetic diversity remains constant across European maize landraces

Maize was likely introduced multiple times to Europe through different routes (Brandenburg et al. 2017). This introduction history and the consecutive expansion should be reflected in the population structure of European traditional varieties. We assessed the population structure within European maize of 1,253 individuals, which were divided into 38 traditional populations and modern elite inbreds of the two heterotic pools, flint and dent, representing a diverse sampling of the European maize (Figure 1a). We performed a PCA based on 491,477 genome-wide SNP markers. The PCA showed the integrity of landrace populations, as individuals from the same landrace grouped together (Figure 1b). The first two principal components explaining 7.1% of the variation, mostly separated the populations into a Western and Eastern cluster.

**Figure 1.**
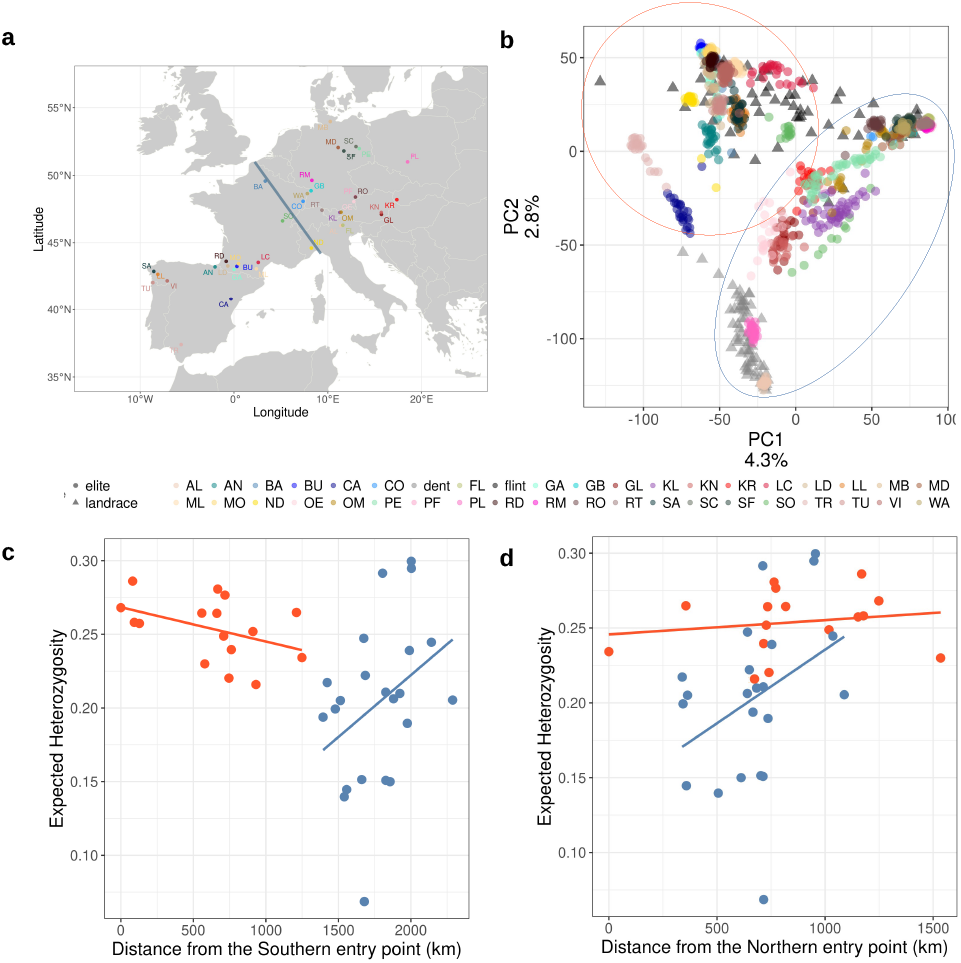
Population diversity of European maize. a. Geographic distribution of the landraces in Europe. Blue line denotes approximate geographic position which separates the populations into two clusters, the Eastern and Western European clusters. b. First two PCs of genomic PCA on 38 landraces and 2 modern elite pools. Color of dots corresponds to population, shape to the type of population; landrace (circle) or elite breeding line (triangle). The two major clusters identified are marked by red (Western Cluster) and blue (Eastern Cluster). b. Mean expected heterozygosity is plotted against the distance from the potential introduction points in Europe via c. the South and d. North Europe. Populations are colored based on cluster membership; Eastern (blue) or Western (red) cluster.

Furthermore, PC2, which explained 2.8% of variation, clearly separated flint (Europe) and dent (US and Europe) maize. It has been shown before that the flint and dent pools are genetically differentiated, despite sharing haplotypes (Brown and Anderson 1947; Unterseer et al. 2016; Haberer et al. 2020). This reflects the long divergent germplasm pools of the two heterotic groups. Brown and Anderson (1947); Unterseer *et al*. (2016). Moreover, there are 2 landrace populations, Polnischer Landmais (PL; Poland) and Altreier (AL; Italy), which cluster with the dent instead of the flint heterotic group. The existence of multiple well defined populations within the European maize landraces was also supported by a STRUCTURE analysis of the individuals (Figure S1). The STRUCTURE supports the presence of 35 (cv = 0.38024) separate populations (Mayer et al. 2017), with varying levels of shared ancestry between the populations.

We estimated genetic diversity using expected heterozygosity (*H*_*Exp*_) per site within each population (Table S1). The median *H*_*Exp*_ was 0.232 and mean *H*_*Exp*_ was 0.224 across all populations (Figure S2a). The highest *H*_*Exp*_ was observed within the landrace Gleisdorfer (GL; Austria) and lowest for Fleimestal (FL; Italy), with *H*_*Exp*_ 0.299 and 0.068, respectively. The elite breeding lines had the highest inbreeding coefficients (median *F*_*IS*_= 0.992), but only a few landraces had high inbreeding coefficients (Figure S2c). Surprisingly, the populations in Eastern Europe showed higher variation in inbreeding than in the West (Figure 1c).

In line with the theory of range expansion, we looked for patterns of decreased *H*_*Exp*_ across the geographic distance from potential introduction points to study the impact of maize range expansion within Europe (Figure 1c, d). We estimated the Haversine geographical distance between the populations closest to the potential entry points of maize in Europe and the rest of the populations. We used the population Tuy (TU) in Spain as the closest population to the entry for introduction via the Southern route and the coordinates of the population Barisis (BA; France) as the closest population to the second entry point in Central Europe, as described in Tenaillon and Charcosset 2011. The *H*_*Exp*_ was significantly negatively correlated with the geographic distance from Tuy (Spain; p = 0.024, *ρ* = -0.366), which might reflect a pattern of declining genetic diversity along the range expansion route. However, the opposite pattern was observed for the Northern route from Barisis (France; p = 0.00099, *ρ* = 0.518). There was a significant negative correlation (p = 0.0191, *ρ* = -0.568) of *H*_*Exp*_ and distance within the Western cluster of populations from the introduction point in the South. No equivalent correlation was detected within the Eastern cluster of populations, or with the introduction point close to Barisis.

### Population structure and expansion across Europe

The calculated *F*_*ST*_ values, which ranged from 0 to 0.531, with the mean and median of 0.255 and 0.257, respectively, support the presence of separate landrace populations (Figure 2a). The Polnischer Landmais (PL; Poland) population, which clustered with the dent elite lines in the PCA, has on average high *F*_*ST*_ values with the other populations (median of 0.352), corroborating the results of the PCA.

**Figure 2.**
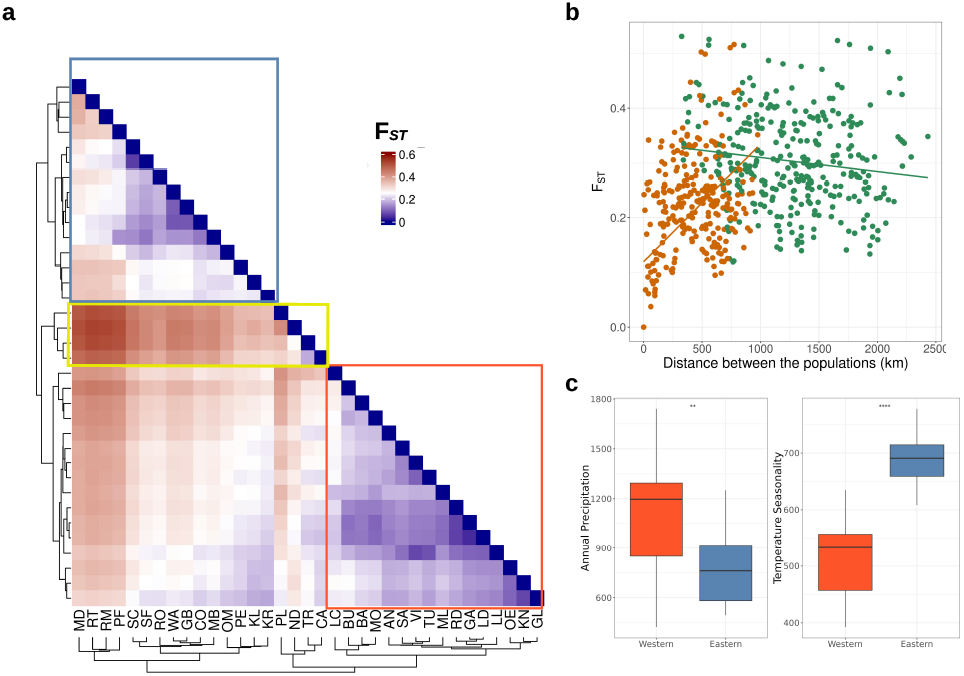
Population clustering of European maize landraces. a. Pairwise *F*_*ST*_ values between 35 European maize landraces. The Western (red box) and Eastern European (blue box) clusters, and ‘dent’ cluster (yellow). b. The *F*_*ST*_ values between each pair of populations against the geographic distance between the populations. Pairs within cluster (either the Western or Eastern European) are colored in orange, pairs belonging to different clusters are shown in green. c. Bioclimatic of landrace geographic origin factors by population cluster (both p<0.05).

The two population clusters, each of which includes landraces from the Western or from Eastern Europe, are also definable by the *F*_*ST*_ values. On average, the *F*_*ST*_ values were significantly lower (Kolmogorov-Smirnov test p = 4.663e-09; Figure S3) between populations within the same cluster (mean of 0.259 and median of 0.2542) than between populations between the two clusters (mean of 0.303 and median of 0.295). A third smaller cluster of populations was observed within the Western cluster, which consists of Polnischer landmais (PL; Poland), Nostrano dell Isola (ND; Italy), Tremesino (TR; Spain) and Castellote (CA; Spain). Those correspond to those landraces, which cluster with the dent elite lines in the PCA (Figure 1b).

We summarized historical bioclimatic variables (Fick and Hijmans 2017) from the origin of landraces, according to the two major clusters we identified in both the PCA and *F*_*ST*_ analysis. The Western cluster is characterized by significantly higher (Wilcoxon’s p = 1.6e-05) annual precipitation and overall higher mean temperature in the coldest quarter and minimum temperatures in the coldest month, with p-values of 9 e-10 and 1 e-06 (Figure 2c). Meanwhile, the Eastern cluster has a significantly (p = 2.6 e-10) greater temperature seasonality than the Western cluster. In general, out of the 19 bioclimatic variables, 13 were significantly different between the two population clusters, when using a p-value cut off of 0.001 (Figure 2c; Figure S3). Therefore, since the population subdivision corresponds well to both climatic and genetic differences, we will use this clustering in the paper in the future.

The genetic differentiation between populations increases with the distance between them (p < 2.2e-16, *ρ* = 0.409; Figure 2b). The same result was reached when we used Slatkin’s linearized *F*_*ST*_ (p < 2.2e-16, *ρ* = 0.409; Figure S4b). The pattern of isolation by distance was also evident when we take into account the environmental distance between the populations instead of the physical one. The correlation between *F*_*ST*_ values and environmental distance was *ρ* = 0.282 (p = 4.882e-13; Figure S4c), with the results being the same when we take into account Slatkin’s linearized *F*_*ST*_. This general pattern of correlation with the geographic distance was stronger within the Western European group (p = 7.14e-16, *ρ* = 0.678), than the Eastern European cluster of populations (p = 1.179e-07, *ρ* = 0.367). This pattern was also observed when the environmental distance was correlate within each cluster, with the Eastern and Western clusters of populations having positive correlations of *ρ* 0.346 (p = 2.5e-10) and 0.471 (p < 2.2e-16), even though the median *F*_*ST*_ value between the Western European populations was 0.167. In contrast, the Eastern European populations had a more defined population structure, with median *F*_*ST*_ of 0.2391.On the contrary, the correlation of *F*_*ST*_ with the environmental distance was higher within each cluster than between clusters. Between the populations of the same cluster the correlation with the environmental distance was *ρ* = 0.290 (p = 9.015e-08), while when the populations belonged in different clusters the *F*_*ST*_ values had a significant negative correlation with the environmental distance (p = 0.0016, *ρ* = -0.179; Figure S4c).

The strongest genetic differentiation was observed between the landrace populations of Rheintaler St. Gallen (RT; Switzerland) and Nostrano dell Isola in Italy (ND; *F*_*ST*_ = 0.531), Rheintaler Monsheim (RM; Germany) and Nostrano dell Isola (ND; *F*_*ST*_ = 0.525), as well as between Rheintaler St. Gallen (RT; Switzerland) and Tremesino in Spain (TR; *F*_*ST*_ = 0.523) despite their geographic proximity to each other (approximately 324km and 559km between RT/ND and RM/ND, respectively). These examples demonstrate the complex local history of maize in Europe.

To further investigate the geographic patterns of diversity, we used the FEEMS analysis, which estimates effective migration surfaces (*λ* = 100; cv error = 0.126; Figure S4d). The effective migration rate was higher than neutral (log10(w) > 1) within the Iberian peninsula, with a barrier to migration present along the Pyrenees (log10(w) <1), reducing the gene flow between the Iberian peninsula and Central Europe. Within the Central European cluster of populations, there were surfaces with lower effective migration (log10(w) <1). Those areas overlap with the locations of the Rheintaler Monsheim (RM; Germany), Nostrano dell Isola (ND; Italy) and Rheintaler St. Gallen (RT; Switzerland), which also have high *F*_*ST*_ values between them (Figure 2c).

### Evidence for selection within two European maize landraces

The observed population structure within Europe could have been the result of selection pressures at the new environment. We scanned the genome of two double haploid (DH) line libraries derived from the Petkuser Ferdinand Rot (PE; Germany) and Kemater Landmais Gelb (KL; Austria) landraces (Mayer *et al*. 2020) for signals of relatively recent positive selection. The iHS statistic measures the relative extended haplotype homozygosity between alleles and indicates signatures of recent selection. Within both Petkuser Ferdinand Rot (PE; Germany) DH and Kemater Landmais Gelb (KL; Austria) DH libraries, we detected SNPs with elevated absolute iHS value (Figure 3).

**Figure 3.**
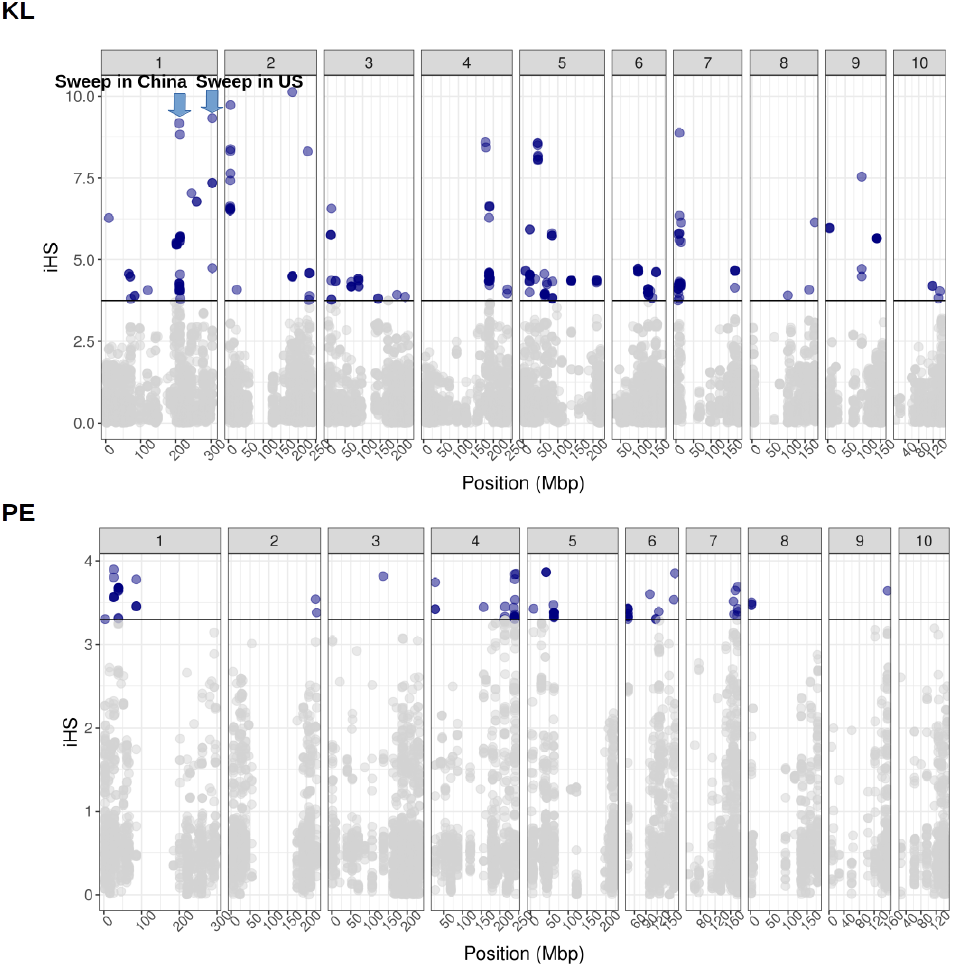
Recent selective sweeps in two European maize double haploid (DH) libraries. The absolute iHS values across the genome are plotted for the Kemater Landmais Gelb (KL; Austria) DH (top) and Petkuser Landmais Rot (PE; Germany) DH (bottom) populations. In blue are highlighted the positions that are significant for being under positive selection (empirical p value cut-off of 0.05). In the Kemater Landmais Gelb population the two locations that have been previously identified as under selection in Chinese or US breeding lines as marked.

In total, we found 266 and 103 significant markers for Kemater Landmais Gelb (KL; Austria) DH and Petkuser Ferdinand Rot (PE; Germany) DH respectively (p-value < 0.05; Table S2). In both double haploid populations the signatures of selection were distributed along all chromosomes. We identified the closest unique genes to the selected SNPs, and specifically, we found 140 genes for Kemater Landmais Gelb (KL; Austria). These were significantly enriched (p < 0.05) for two Gene Ontology functions ‘lipid metabolic process’ and ‘cellular lipid metabolic process’ (Figure S5; Table S2). Moreover, the markers on chromosome 1 overlapped with previously identified selective sweep regions related to maize breeding in the US and China (Huang et al. 2022). Only 48 unique genes were located in close distance upstream or downstream from the identified selection signals in the Petkuser Landmais Rot (PE; Germany) DH, but we did not find overlapping regions under selection in the two populations. Those genes were significantly enriched for the 18 GOs (p < 0.05), including as the top 3 the functions “regulation of nitrogen compound metabolic process”, “regulation of cellular process” and “regulation of biological process” (Figure S5; Table S3). Hence, this might reflect selection based on the local environment rather than the introduction to Europe or the general breeding process.

### Climatic gradients along the expansion routes could explain part of the observed genetic variance

We further investigated the link between the local environment and genetic diversity within European maize landraces using a redundancy analysis (Capblancq and Forester 2021). We used the historical bioclimatic factors for this purpose, latitude and longitude, and corrected for population structure by using neutral variation. We identified three bioclimatic factors, which explained the largest variance. The first bioclimatic variable was “mean temperature of the driest quarter” (adjusted *R*^2^ = 0.132; p = 0.002), then followed the “precipitation seasonality” (adjusted *R*^2^ = 0.159; p = 0.028) and “precipitation of coldest quarter” (adjusted *R*^2^ = 0.176). Interestingly, two of those bioclimatic variables are also significantly different between the Eastern and Western population clusters. The “mean temperature of driest quarter” and the “precipitation of the coldest quarter” are significantly higher in the environment of the Western cluster than the Eastern one, with Wilcoxon test p values of 3.6e-05 and 4.8e-05, respectively.

When we correlated each population’s value for the three bioclimatic variables with the distance to the two different hypothetical introduction points of maize in Europe, we were able to identify gradients of climatic change across the expansion routes. Along the introduction route starting in Spain, the bio-climatic variables “mean temperature of driest quarter” and the “precipitation of the coldest quarter” were significantly negatively correlated (p < 2.2e-16), with *ρ* values of -0.76 and -0.74, respectively. On the contrary, the bioclimatic variable “precipitation seasonality” had a significant positive correlation of *ρ* = 0.476 (p = 0.0027) with the distance from the introduction point within the Eastern Cluster in Barisis (BA; France). However, the RDA model that had the best fit on the dataset, based on the adjusted *R*^2^ value of 0.39 (p = 0.001), was the full model, which incorporates information about the latitude, longitude and population structure besides the three bioclimatic variables (Figure S6). Therefore, we can infer that the climatic conditions along the expansion routes have partly contributed to the genetic diversity across the European landscape.

### Change of genetic load along the introduction routes and due to breeding

Range expansion has been previously associated with the accumulation of deleterious mutations and overall genetic load (de Pedro *et al*. 2021; Peischl *et al*. 2013). We estimated the genetic load per population, including the elite breeding pools, using the GERP scores from Kistler *et al*. (2018). Estimates of genetic load per population differed significantly (p < 2.2e-16), with the population Rheintaler Monsheim (RM; Germany) having the highest median genetic load (sum of GERPs = 29,881.23), while the population with the least accumulated genetic load (sum of GERPs = 29,578.20) was Bugard (BU; France) (Figure S7). We further tested for correlation between accumulated genetic load and the distance from the two hypothetical introduction points in Southern and Central Europe and found significant correlations of *ρ* = -0.16 (p = 1.18e-11) and *ρ* = -0.156 (p = 3.845e-11), respectively. The accumulated genetic load also had a significant (p < 2.2e-16) negative correlation with the bioclimatic factor “mean temperature of the driest quarter” of *ρ* = -0.214. However, there was no positive correlation with the other two bioclimatic factors identified as important for partitioning the dataset’s variance.

Furthermore, we tested the differences in genetic load between landrace and elite groups, as inbreeding, drift and selection during breeding might have changed genetic load. We grouped both the landraces and the elite lines based on their germplasm origin of either dent or flint as identified in the PCA (Figure 4a; Table S1). Genetic load was significantly higher in both elite pools than their counterpart landrace group, with p-values of 0.0486 and 0.01346 for flint and dent germplasm respectively. The accumulating genetic load was also different between the dent landrace germplasm lines and the flint elite germplasm lines (p = 0.0162). Those differences were not observed for highly deleterious mutations. Both elite germplasm pools had a significantly (p < 2.2e-16) lower number of highly deleterious mutations than the landraces (Figure 4b), suggesting potential purging during breeding. In contrast, fixed load was significantly higher (p < 2.2e-16) in the elite lines than the landraces (Figure 2c), while the segregating load was significantly higher (p < 2.2e-16) in the landraces (Figure 2d). Those differences were observed irrespective of whether the lines belonged in the flint or dent germplasm.

**Figure 4.**
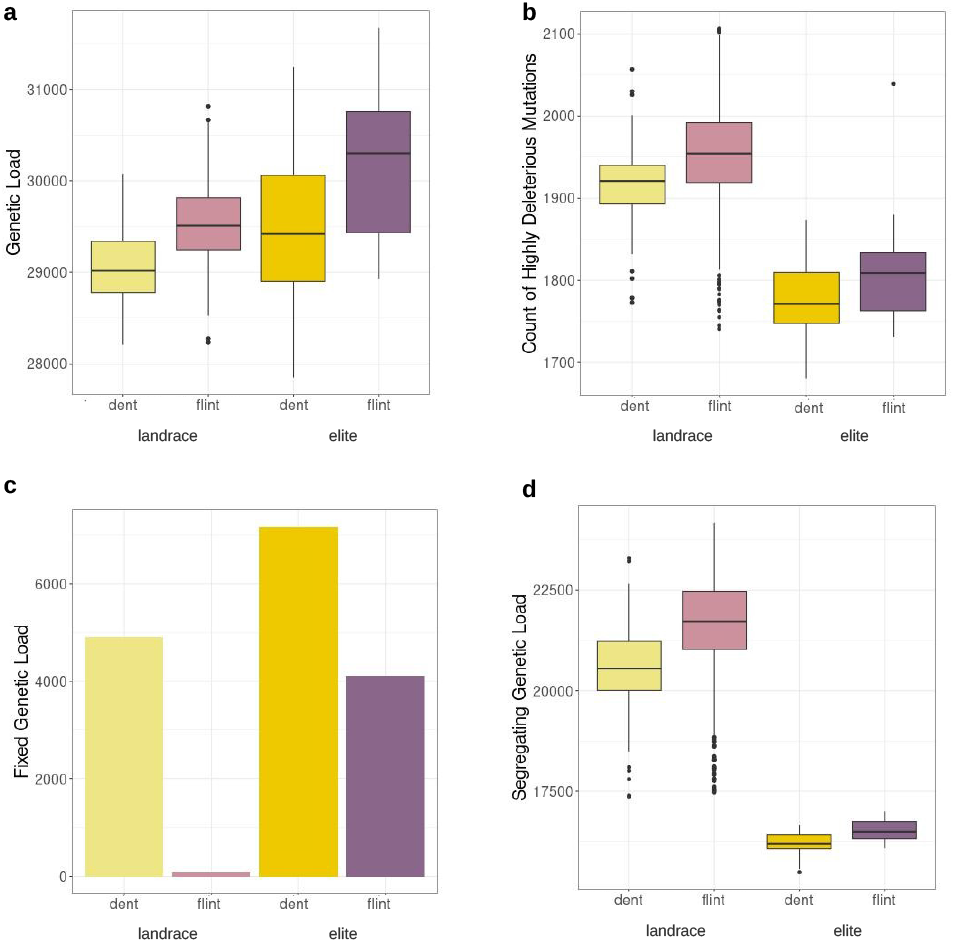
Purging segregating deleterious genetic load within the elite breeding lines. a. Total genetic load estimated, as the sum of the GERP scores for all sites within an individual for dent landraces, flint landraces, flint elite lines and dent elite lines. b. Number of highly deleterious alleles. Count of derived alleles at sites with GERP score larger than 5. c. Fixed genetic load for each group, given as the sum of the GERP scores at fixed derived alleles within within each group of maize lines. d. Segregating genetic load per individual for each of the four groups as the sum of GERP scores of the segregating sites.

## Discussion

### Genetic diversity is shaped by local factors rather than the introduction route

Maize has been introduced to Europe (Janick and Caneva 2005) within the last 500 years and its range expanded rapidly throughout the European continent. Evidence exists for two introductory routes to Europe; one from the Caribbean via Southern Europe and one to Central Europe with material from North America (Tenaillon and Charcosset 2011; Brandenburg et al. 2017). Along this expansion route, local openly pollinating traditional varieties have formed and have been kept and traded by local farmers for centuries. Indeed, the population structure and relatedness between samples of these traditional varieties trace the history of maize in Europe. We used a large sample of European maize landraces to explore how the introduction and expansion of maize in Europe has shaped genetic structure and diversity. The wide geographic sampling and representation of each lan-drace population enabled us to track the impact of colonization of novel environments in a more detailed scale. The analysis of genetic diversity within European landrace populations showed three major clusters across populations (Figure 1). One cluster, which includes dent elite lines, shows that 4 landraces are most likely derived from North American dent populations, while all other landraces clustered with flint elite lines, confirming their flint ancestry and their involvement in the creation of European maize hybrid pools (Mayer *et al*. 2020; Tenaillon and Charcosset 2011; Brandenburg *et al*. 2017). The four landraces have been previously noted for their difference in genetic composition from the rest of the European landraces (Mayer *et al*. 2017). The further separation into two groups within the flint cluster across Europe might represent the two introduction routes of maize into Europe (Tenaillon and Charcosset 2011).

Range expansion can have a strong influence on the genetic diversity of the expanding species, specifically by decreasing genetic diversity and increasing genetic differentiation between the newly established and core populations (Excoffier *et al*. 2009). Despite the pattern of introduction routes and ancestry within Europe, we did not observe a pattern of decreasing genetic diversity along the potential introduction routes. We used the potential introduction locations of maize from Brandenburg *et al*. (2017) to investigate the impact of expansion along the separate routes, but did not find a pattern that would be expected for the colonization by a natural species (Zimmermann *et al*. 2014; Smith *et al*. 2020), which indicates the maintenance of genetic diversity across the continent. On the contrary, genetic diversity is maintained in similar levels across the studied maize populations, suggesting a stronger influence of local maintenance and potential adaptation. This is reflected in the phenotypic level; some of the European maize landraces have been documented to show differences in early vigor, plant height and tillering (Hölker *et al*. 2019), as well as flowering (Balconi *et al*. 2024). However, part of this pattern can be explained by the presence of two distinct germplasm pools. Within Europe, the observed pattern is different from what has been observed within the Americas, where a decrease in genetic diversity has been documented (Huang *et al*. 2022). American maize landraces have followed an expansion route from Central Mexico to North and South America, allowing tracking of changes in genetic composition along the route (Tenaillon and Charcosset 2011; Huang *et al*. 2022; Yang *et al*. 2023). In Europe, however, the pattern is less clear and suggests long range dispersal and a more rapid spread through the continent (Figure 1 and Figure 2).

In potato and European bean, admixture and different admixture histories in the introduced range have been suggested as reason for maintenance of diversity in introduced ranges but often little population structure is observed within a smaller geographic region, such as in the Nordic countries (Ortiz *et al*. 2023; Gutaker *et al*. 2019; Ames and Spooner 2008; Bellucci *et al*. 2023). Admixture between Central and Southern European maize populations might be prevalent (Bradbury *et al*. 2007; Mayer *et al*. 2017; Unterseer *et al*. 2016), however, we find that not only there is population structure dividing the landraces into multiple geographic distinct populations, but also there is an evident pattern of isolation by distance. We identify smaller local genetic clusters between landraces, including one along the Rhine and one into northern Germany that consisted of four populations (Figure 2). The *F*_*ST*_ values between those populations and their neighboring ones are up to 0.5. This sharp distinction likely represents historical trade routes and political and linguistic borders in the area during introduction of maize to Europe. Tenaillon and Char-cosset (2011) described that Caribbean maize was introduced to Southern Europe by Columbus in the 15th Century and from there it was introduced to the Vatican and Italy.

### Genomic regions selected during maize introduction due to adaptation to local environments

We used two very large populations of DH lines derived from divergent landraces to infer signals of recent selection in the landraces. The advantage of such a population is their natural phasing through the DH process, which removes problems with computational phasing in highly heterozygous outcrossing maize. We were able to detect signals of positive selection within the Central European Petkuser Ferdinard Rot (PE; Germany) landrace and Kemater Landmais Rot (KL; Austria) landrace. We found 53 and 3 potentially selected loci in each population but no overlap in selection signals, confirming the importance of local selection in the formation of these local landraces. In addition, we found a partial overlap of detected selection signals in one of the two populations with selective sweeps related to breeding efforts in American and Asian breeding maize lines (Huang *et al*. 2022), suggesting convergent selection targets in different regions of the world. Additional selection signals might be linked to adaptation to the novel environments and subsequent breeding targets that maize encountered after its arrival in Europe. Compared to the Caribbean and Mexican environments, which range from tropical highlands to wet and dry lowlands, temperate European environment offered novel challenges to adapt to (Bellon et al. 2011; Brandenburg et al. 2017). Even though we have only tested selection in two populations, the traces of selection corroborate with the results of the population structure within European maize. Local adaptation can be achieved by even small changes in allele frequencies within populations (Le Corre and Kremer 2012), especially for polygenic traits such as the ones observed here. Regions of high genetic differentiation have been found to harbor genetic variation linked to adaptation to local climate in various plant species (De La Torre et al. 2019) and it can occur rapidly during range expansion (Colautti and Barrett 2013), even if there is extensive gene flow (Hämälä and Savolainen 2019).

The local environment can have profound effects on genetic diversity. Patterns of maintained genetic diversity and population structure in the short evolutionary times of European maize might be related to local environmental adaptation, but they have also been connected to human activity in other crops (Bellucci et al. 2023). For instance, human activity has facilitated the geographic spread and genetic differentiation of rice (Courtois et al. 2012). Within Southern Europe, landrace variability and genetic structure of tomato landraces, which were introduced from South America at a similar time as maize, are related to the fruit type (Corrado et al. 2014; García-Martínez et al. 2013), a highly coveted breeding trait.

We document three historical bioclimatic factors, mean “temperature of the driest quarter”, “precipitation seasonality” and “precipitation of coldest quarter”, which significantly explain part of the genetic variance. This indicates that at least part of the genetic variation is due to adaptation to the more diverse environments of the Iberian peninsula and Central Europe. The idea is reinforced both by the significant differences of the two European clusters and also by the gradual change of the significant bioclimatic factors along the possible introduction routes in Europe. Historically, crops such as maize and potato were introduced in Europe during the 15th to 17th centuries to solve problems created by famines, which took place due to cold and dry years (Ljungqvist *et al*. 2024). The three historical bioclimatic factors could be related to the tendency of farmers to select traits that would improve the survivability of maize plants under those conditions in which wheat failed.

However, we cannot exclude the possibility that some traits were selected as reaction to local micro-environments. For instance, within the double haploid population of Kemater Landmais (KL; Austria), genes close to the markers under selection are enriched for the function ‘lipid metabolic process’ and ‘cellular lipid metabolic process’. Tropical maize lines have adapted to colder temperatures via introgression of alleles related to lipid metabolism (Barnes *et al*. 2022). The Petkuser Ferdinard Rot (PE; Germany) population showed enrichment for functions such as “obsolete positive regulation of nitrogen compound metabolic process”. Adaptations to low nitrogen in sub-Saharan African maize has been documented (Worku *et al*. 2007).

### Local purging of genetic load due to inbreeding and selection

Finally, we estimated the genetic load within the European landrace populations. In maize, the deleterious mutations accumulate within complex phenotypes of interest, even though, in overall, they are maintained in low frequencies (Mezmouk and Ross-Ibarra 2014). However, accumulation of genetic load during range expansion has been observed in the Americas (Wang *et al*. 2017). Here, we observed that genetic load within the European landrace populations is variable but does not follow a strong pattern of accumulation in any cluster. Population sub-division, similar to the one observed here, could maintain the genetic load in similar levels (Glémin *et al*. 2003), though the observed migration between the different landraces could have also helped to alleviate the accumulation of load across the expansion route (Theodorou and Couvet 2006). Additionally, the introduction of maize in Europe has been relatively recent, and therefore the crop spread throughout the European continent relative fast. This fast range expansion could have limited genetic load accumulation (Gilbert *et al*. 2018), leading to the absence of a correlation between load and geographic distance from the possible introduction points in Europe. On the other hand, the adaptation to climatic parameters and environmental gradient can promote the accumulation of genetic load (Gilbert *et al*. 2017; Fiscus *et al*. 2024), maintaining differences between populations as we observe here. Indeed, we observe a significant correlation between one of the bioclimatic factors (mean temperature during of the driest quarter) with the accumulation of genetic load. This bioclimatic parameter is significantly associated with differences between the two population clusters and has a positive association with longitude. Together, they add support to the partial impact of environment to accumulation of load, but the correlation between location and environment might obscure the causal reason for load accumulation.

When we compared the landraces to the elite breeding lines in our populations, we found that the number of highly deleterious mutations decreased in the elite lines, while fixed load was significantly higher (Figure 4b and 4c). These results suggest purging of highly deleterious alleles but fixation of mildly deleterious load, potentially due to inbreeding and decreased population size during selective breeding. For example in *Arabis alpina*, selfing in contrast to outcrossing has led to a decrease in the highly deleterious load (Zeitler *et al*. 2020). Inbreeding can effectively remove the deleterious mutations particularly when they are recessive, making it more effective than selection alone (Glémin 2003), which might explain the higher load in landrace populations. Purging of highly deleterious mutations has been shown to decrease load in maize compared to teosinte, reducing inbreeding depression in maize compared to teosinte (Samayoa et al. 2021). Moreover, the need for inbred fitness in breeding programs would further favor purging during breeding and can lead to the removal of especially lethal mutations by inbreeding (Wang et al. 1999). Yet, the high genetic diversity observed within the landraces might enable masking of deleterious alleles but leads to the higher segregating genetic load. Masking of fixed mildly deleterious alleles in hybrids might contribute to heterosis in European maize hybrids (Yang et al. 2017a), but further reduction of fixed genetic load might enable further selective gain.

Taken together, our results illustrate the complexity of the introduction of maize to Europe and the individual history of landraces within the continent and provide an example of local populations maintaining a broad genetic diversity. We have expanded on the knowledge on the genetic composition of the European landraces by exploring how it was shaped by possible introduction events, adaptation to novel environments, as well as drawing an example of load accumulation in a rapid range expansion and the impact of breeding on rare alleles.

## Data Availability

All data used are publicly available from their respective publications of origin (Unterseer *et al*. 2016; Mayer *et al*. 2017, 2020). The merged dataset will be made available upon publication. All scripts used in the analysis are available under https://github.com/cropevolution/europeanMaize.git.

## Author contribution

MGS conceived the study. MT and KS conducted the analysis. MT prepared figures and tables. MT, MGS and KS wrote the manuscript. All authors discussed the results, edited, and approved the manuscript.

## Acknowledgments

We would like to thank A. Singh for valuable feedback on the manuscript. We acknowledge funding by the Deutsche Forschungsgemeinschaft (DFG, German Research Foundation) under Germany’s Excellence Strategy – EXC-2048/1 – project ID 390686111 and financial support by the Federal Ministry of Education and Research (BMBF, Germany) under the Plant Breeding Research for the Bioeconomy initiative (funding ID: 031B0195, project MAZE).

## Supplement

**Figure S1.**
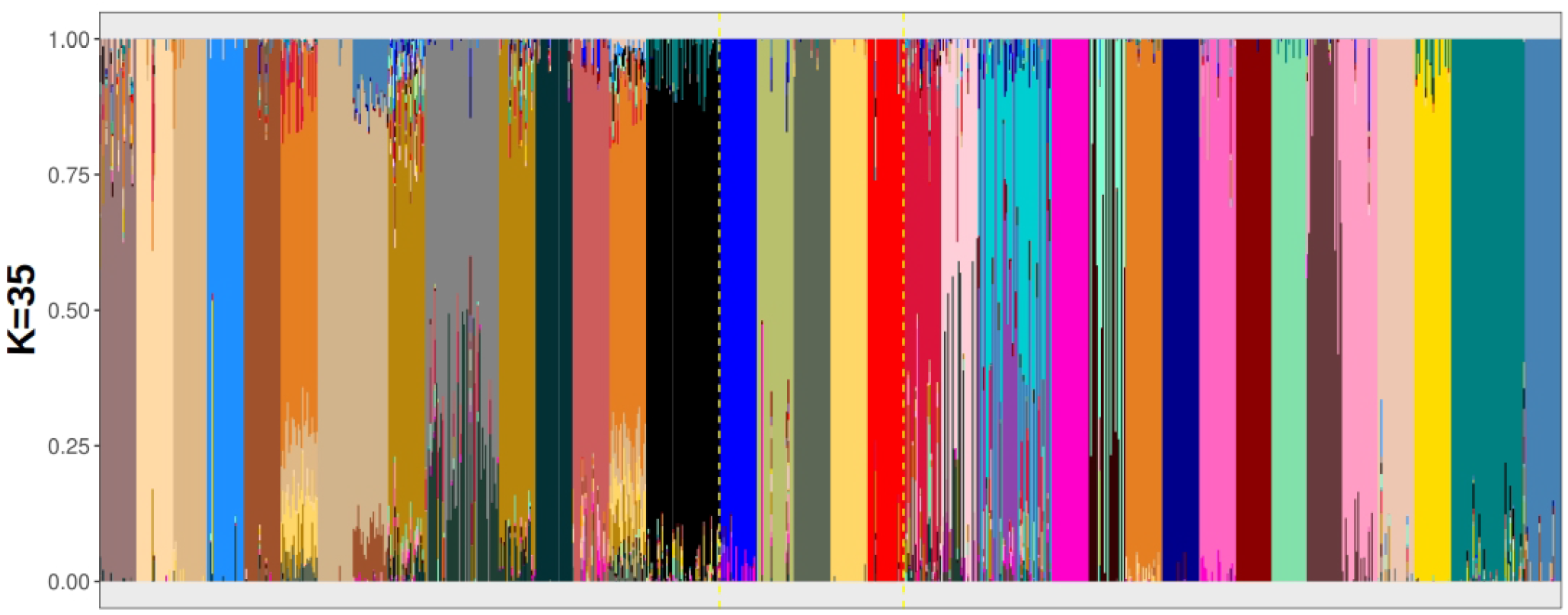
Genetic diversity within European maize. a. The density of the mean *H*_*Exp*_ for all populations is shown. b. The density of the *F*_*IS*_ values for each individual of all landraces and elite lines. c. The *F*_*IS*_ values per population, ordered from West to East with the elites depicted at the far right of the plot.

**Figure S2.**
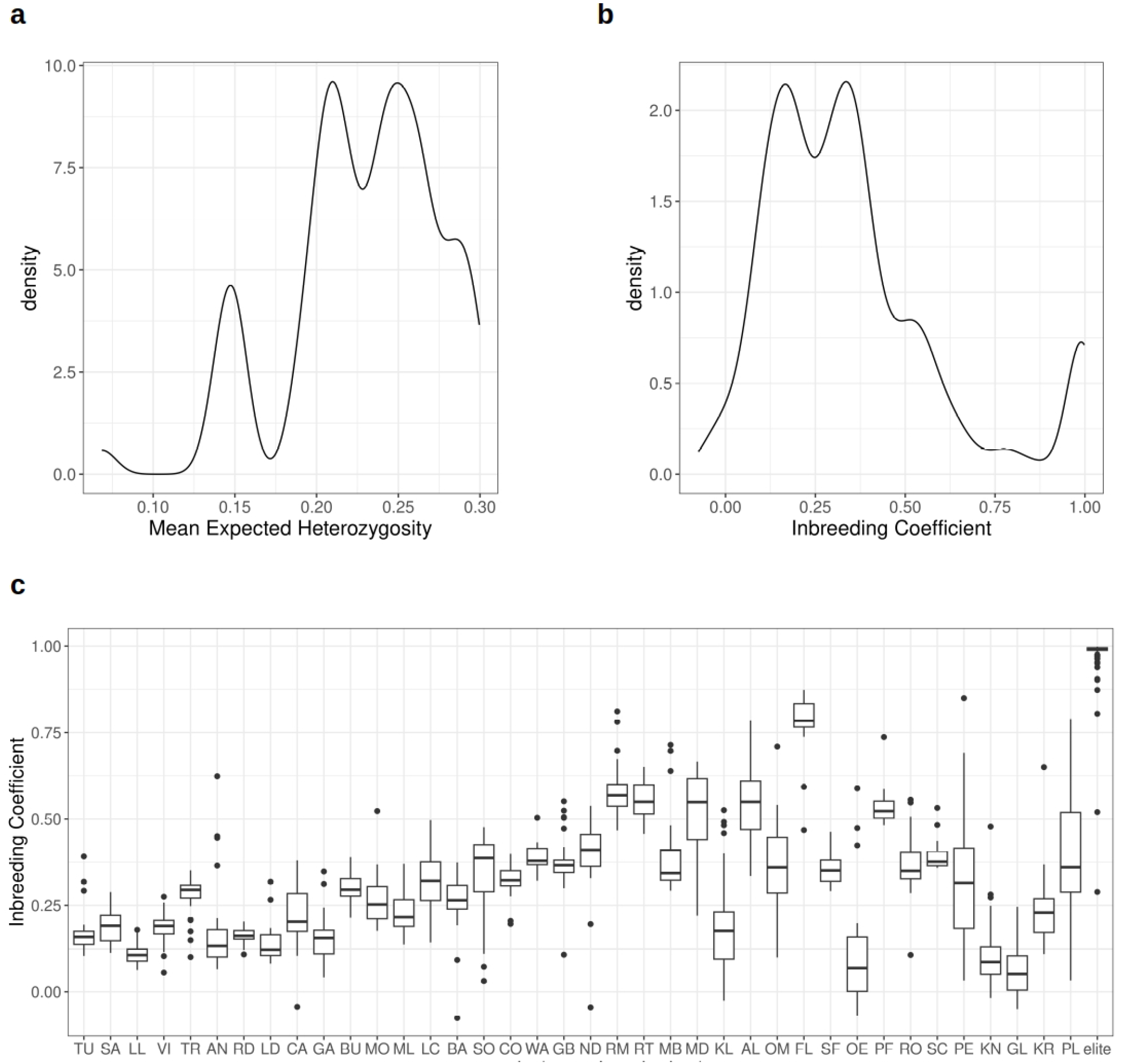
Structure analysis of the European maize landraces. Result for K=35, which was the optimal number of clusters are shown. The populations are ordered as in the *F*_*ST*_ heatmap (Figure 2). The dashed yellow lines in both plots indicate where each *F*_*ST*_ group starts and ends.

**Figure S3.**
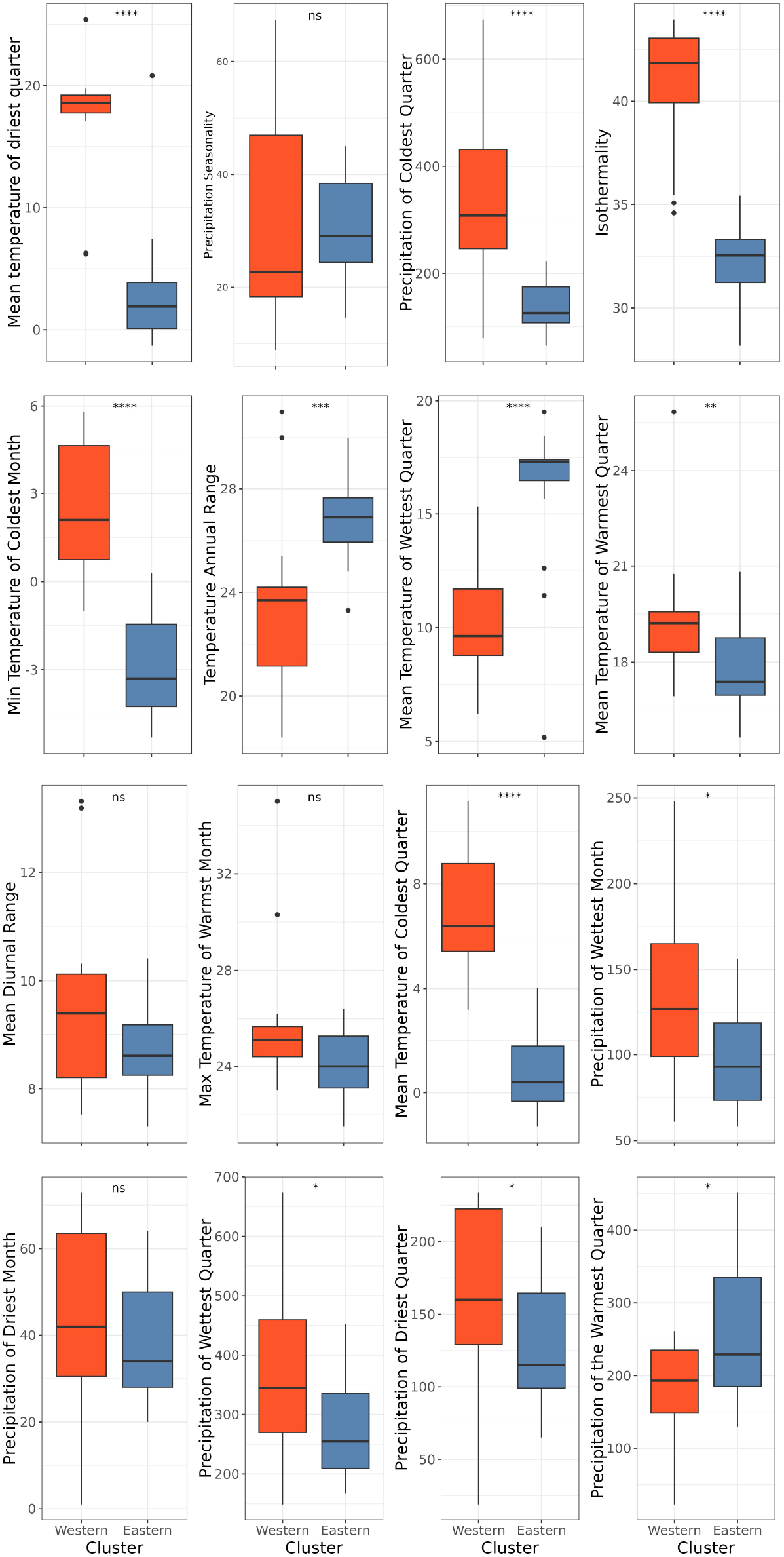
Environmental diversity between the Eastern and Western European population clusters, comparing the historical bioclimatic factors bio1-bio19. The significance level for each parameter is indicated with ‘ns’ for p > 0.05, * for p <=0.05, ** for p<=0.01, *** for p <=0.001 and **** for p<=0.0001.

**Figure S4.**
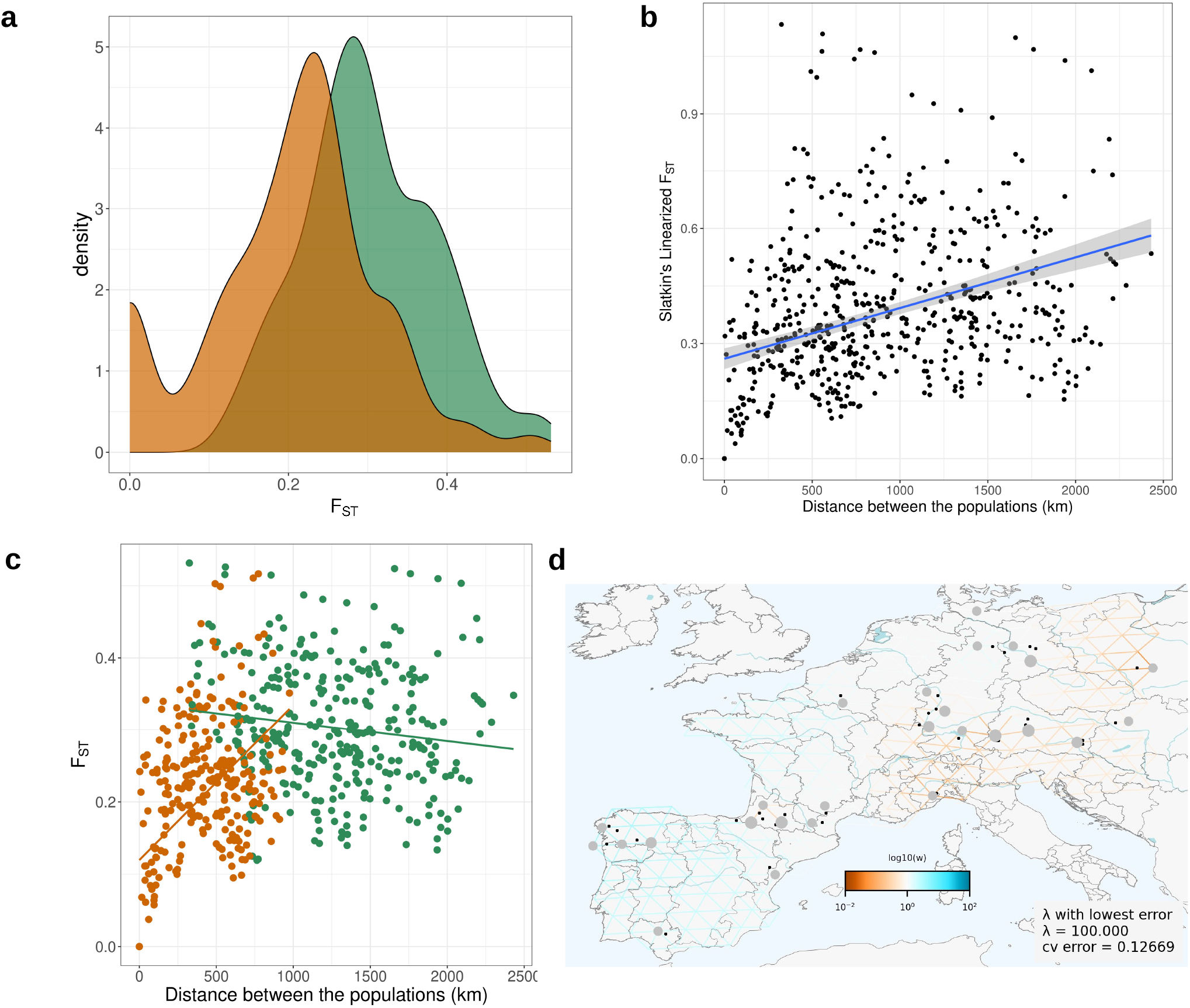
a. The distributions of pairwise *F*_*ST*_ values for all the pairs within the same cluster (orange), or between the two clusters (green). b. Slatkin’s Linearized *F*_*ST*_ against the distance between the populations in kilometers. c. *F*_*ST*_ values are plotted against the environmental distance between the populations. The values in green show pairs of populations belonging to different clusters and orange pairs of populations belonging to different clusters. d. Effective migration rate within Europe. The grid is colored blue when estimated effective migration rate lower than expected and red when it is higher than expected.

**Figure S5.**
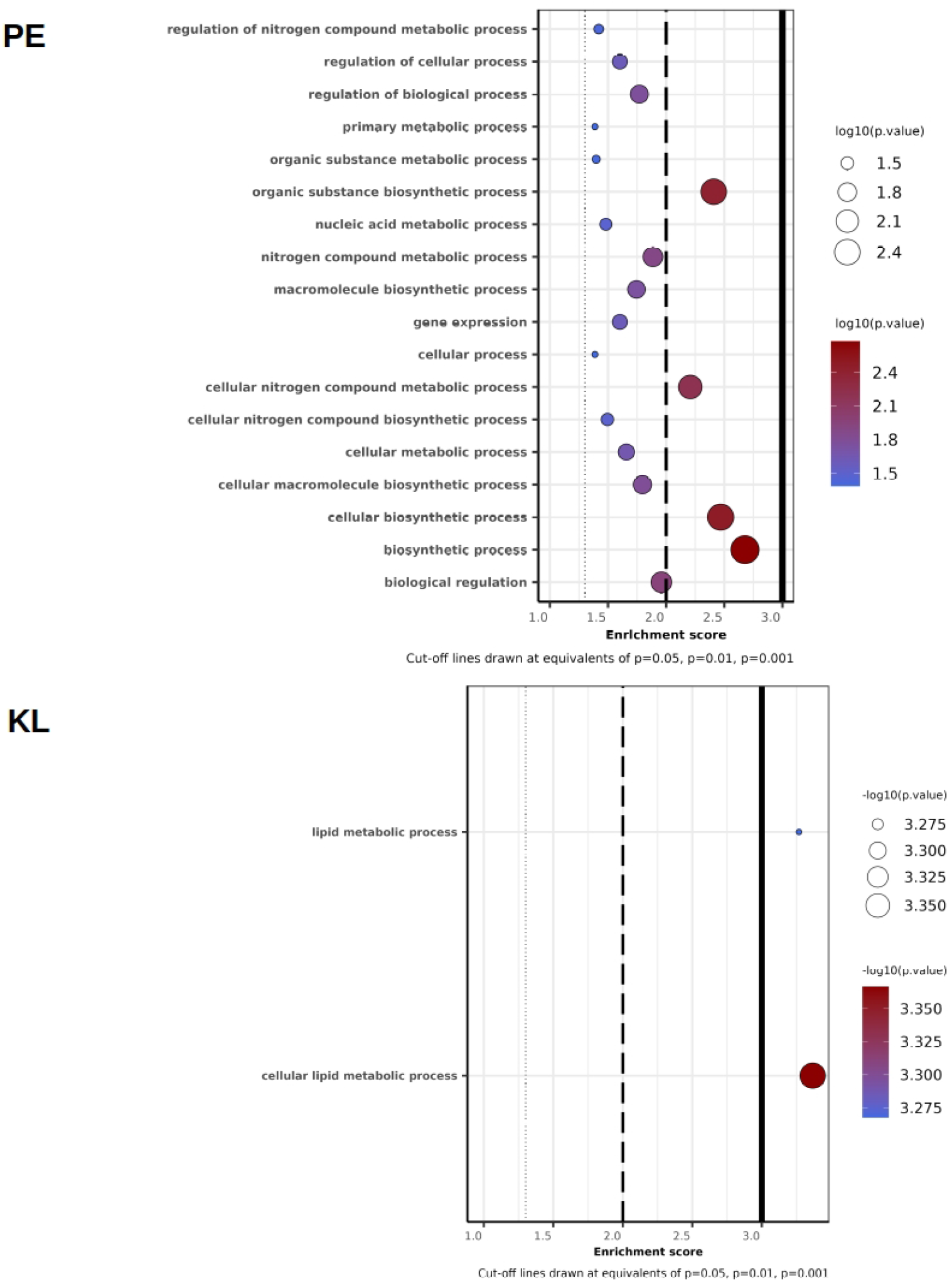
Enriched Gene Ontology categories for the genes closest to the markers with selection signal for Petkuser Ferdinard Rot DH lines (PE) on the top and Kemater Landmais Gelb DH lines (KL) on the bottom.

**Figure S6.**
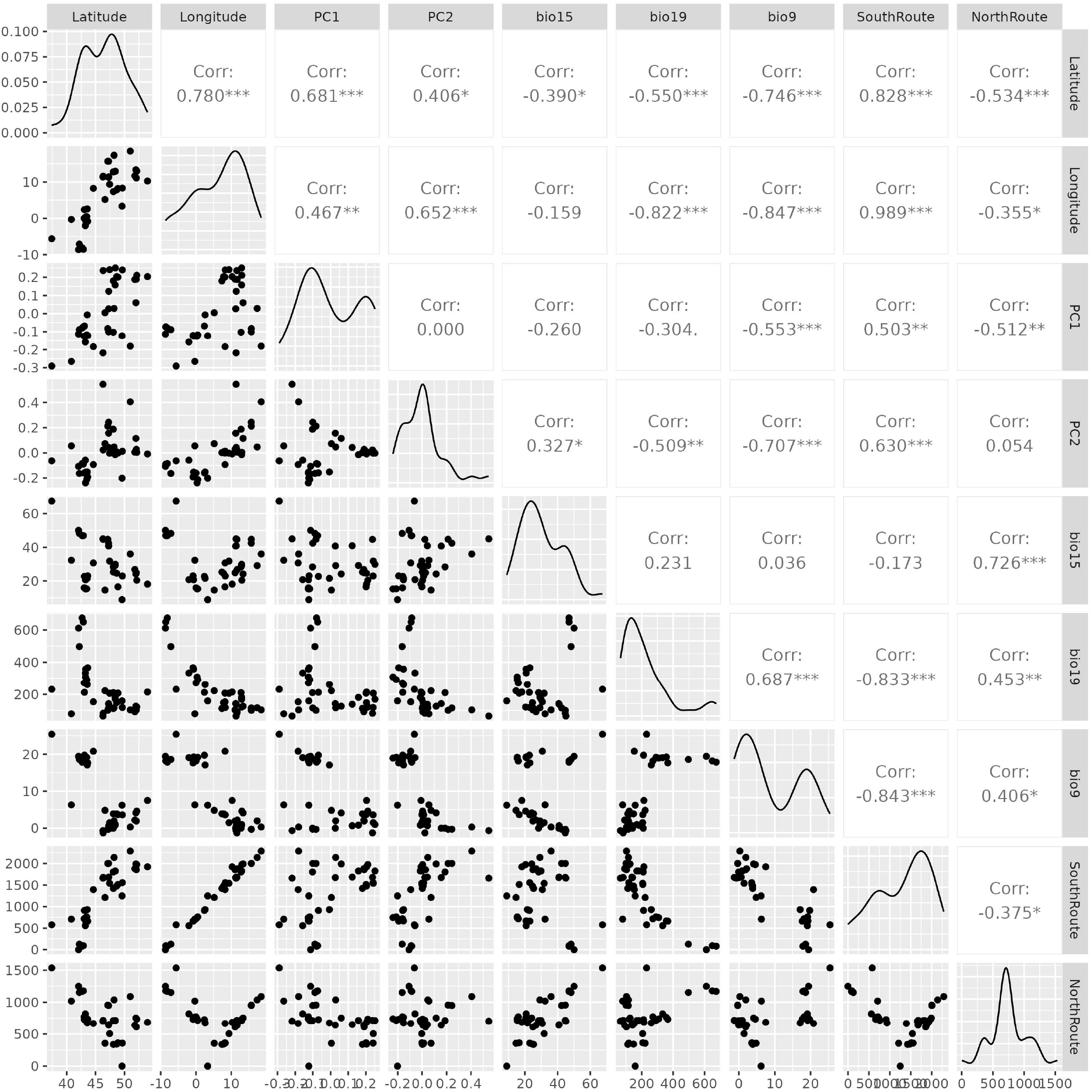
Environmental and Genetic diversity parameters explaining genetic variation based on a redundancy analysis.

**Figure S7.**
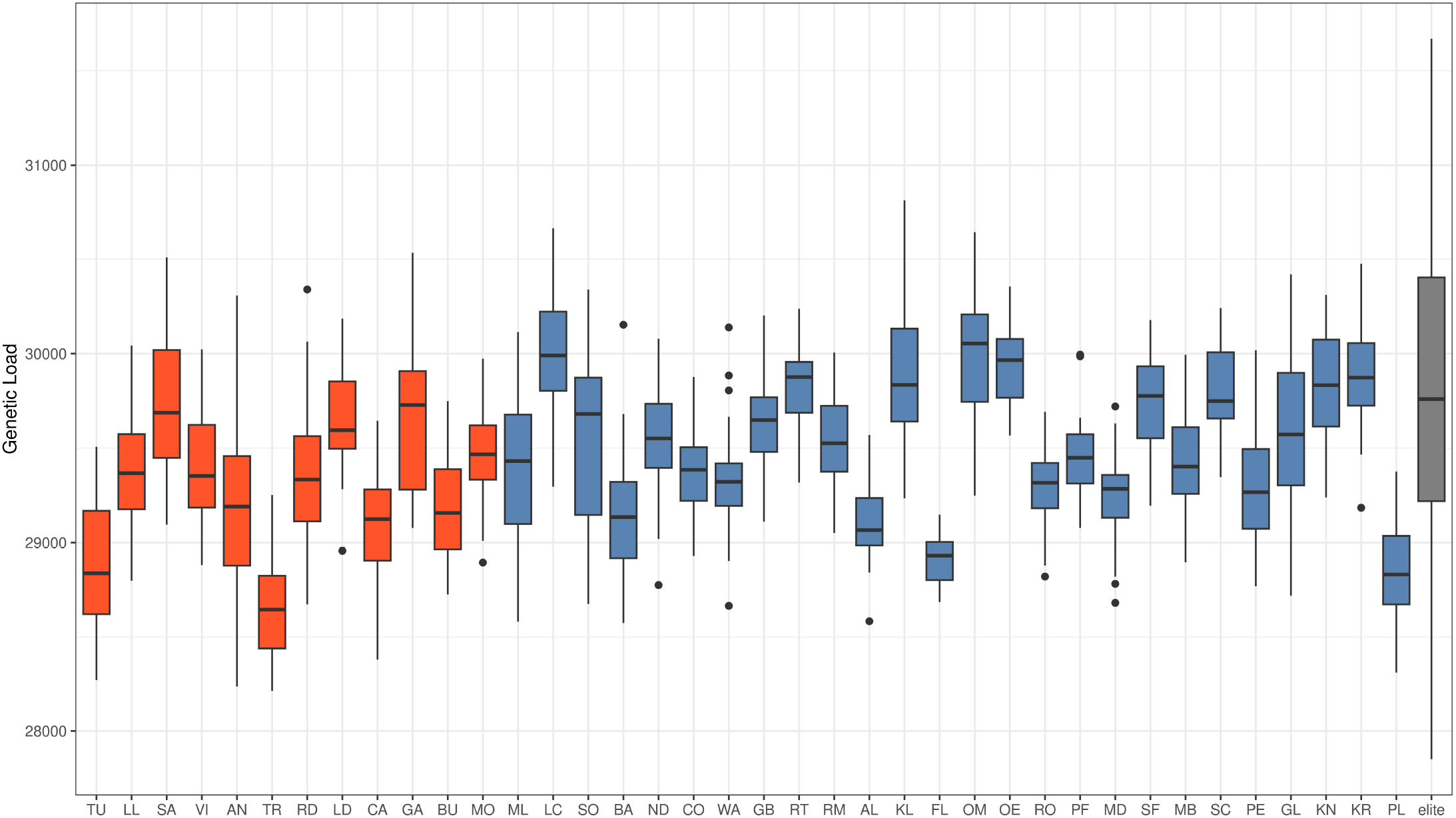
Genetic load per European landrace population ordered by their distance from the hypothetical Southern introduction point. In red, all populations belonging in the Western cluster, in blue populations of the Eastern cluster and in grey the elite lines.

